# Discernibility in explanations: an approach to designing more acceptable and meaningful machine learning models for medicine

**DOI:** 10.1101/2025.04.06.647498

**Authors:** Haomiao Wang, Julien Aligon, Julien May, Emmanuel Doumard, Nicolas Labroche, Cyrille Delpierre, Chantal Soulé-Dupuy, Louis Casteilla, Valérie Planat-Benard, Paul Monsarrat

## Abstract

**Background:** Although the benefits of machine learning (ML) are undeniable in health-care, explainability plays a vital role in improving transparency and understanding the most decisive and persuasive variables for prediction. The challenge is to identify explanations that make sense to the biomedical expert. This work proposes discernibility as a new approach to faithfully reflect human cognition, with the user’s perception of a relationship between explanations and data for a given variable.

**Methods:** A total of 50 participants (19 biomedical and 31 data scientists) evaluated their perception of the discernibility of explanations from both synthetic and human-based dataset (National Health and Nutrition Examination Survey). The inter-rater reliability was tested through the intraclass correlation coefficient (ICC). 13 statistical coefficients were considered to be able to capture for a given variable the relationship between its values and its explanations. A Passing-Bablok regression was performed for each user to highlight the consistency between user rating and each coefficient.

**Findings:** The low inter-rater reliability of discernibility (ICC¡0.5) with no difference between areas of expertise or level of education underlines the need for an objective metric of discernibility. Among all evaluated metrics, *dcor* metric was found to be the most suitable to capture the intra-individual reliability of discernibility perceived by users (median slope closer to 1 and a narrower confidence interval width for the Passing-Bablok regression with the lowest differential bias between the most and least discernible values).

**Interpretation:** *dcor* was shown to be a reliable metric for assessing the discernibility of explanations, effectively capturing the clarity of the relationship between the data and their explanations, and providing clues to the underlying pathophysiological mechanisms that are not immediately apparent when examining individual predictors. Discernibility can also serve as an evaluation metric for model quality, used to prevent overfitting or aid in feature selection, providing medical practitioners with more accurate and persuasive results.

## 1. Introduction

Although the benefits of data exploration and analysis models based on machine learning (ML) are undeniable in many fields, such algorithms have become increasingly complex and poorly understandable by humans [1], hence considered as “black-boxes”. In healthcare, where each decision directly impacts patients’ life, medical practitioners are concerned about the application of ML models due to this lack of transparency [2]. This issue has also been echoed by government agencies and international organizations in the form of artificial intelligence (AI) development guidelines that extend to regulations. European Union’s *AI Act* ^1^ *categorizes AI systems that pose significant risks to individual health as ‘high risk’, mandating transparency to counteract the potential incomprehensibility or complexity of these systems. In the United States, Executive Order on the Safe, Secure, and Trustworthy Development and Use of Artificial Intelligence* ^2^ emphasizes protecting patients, and encourages the regulatory agencies to delineate clear requirements and expectations related to the transparency of AI models. Explainability in AI (XAI) plays a vital role in improving transparency in the ML process by helping “users” to understand the predictions made by models, what are the most decisive variables in prediction, whether at the level of groups of patients and/or at the individual level [3]. This is crucial for a biomedical use, in advancing patients’ rights to information and autonomy, ultimately strengthening the relationship between patients and medical practitioners [4, 5]. This is all the more important as this explainability can also lead to a better understanding of pathophysiology and the underlying biological mechanisms, even if the inference of causality must be carefully considered.

Among all XAI methods, feature importance-Based XAI methods, such as Local Interpretable Model-Agnostic Explanations (LIME)[6] and SHapley Additive exPlanations (SHAP)[7], have received substantial interest from both XAI and biomedical researchers [8, 9]. These methods facilitate the investigation of each single individual, feature by feature, assist in identifying compelling individual patient cases and noteworthy features. Despite the increasing use of XAI, a consensus of a “good explanation” has yet to be established. Researchers have often defined multiple metrics to assess XAI from various points of view. Doshi-Velez and al. [10] proposed a three-level taxonomy of evaluation approaches, classified as functionally grounded (based on a formal definition), or human-based and application-based evaluations, which are human-centered evaluations. As XAI is an inherently human-centered approach [11], human involvement is crucial in evaluations, while the main drawback is the cost of these manual evaluations (e.g. the recruitment of participants, the time and the sophisticated design to alleviate cognitive bias). In contrast, the functionally-grounded evaluations are less cost-intensive, but concentrate on objective functionality, such as correctness (faithfulness of the explanation with the predictive model), consistency (identical inputs have identical explanations), and compactness (size of the explanation) [12].

This work proposes *discernibility* and its associated statistical metric as a new concept for objectively evaluating explanations while reflecting human cognition. It takes into account, at the same time, the effectiveness of functionally-grounded evaluation, and the human factor of human-involved evaluations. Inspired from the term *readability* [13], *discernibility* faithfully reflects the user’s perception of a relationship between explanations and data for a given variable. This work also demonstrates the usefulness of *discernibility* as a complement to model performance in feature selection, providing accurate but also more persuasive results to medical practitioners.

## 2. Methods

### 2.1. Study design

STARD 2015 guidelines [14] were followed. The study was conducted from November to December 2023. The aim was to assess the consistency between users’ perceptions of the discernibility of XAI explanations and established statistical and objective metrics. As there is no universally recognized standard, the initial phase of this study involved assessing a representative set of explanations by the users. Real and synthetic explanations were included to ensure completeness. Participants were instructed to evaluate all explanations in randomized order and were kept blind to the scores tested during the study to mitigate potential order bias and anchoring effect.

### 2.2. Participants

Since users from different academic background might have different explanation preferences [15], the study has included users from biological/medical or data science (including XAI) area of expertise, with or without already XAI experience and with different level of education. Both factors were used for subgroup analyses. Participants were recruited from a biological research institution (RESTORE lab, Toulouse, France) with both physiologists and physicians, a computer science research institution (IRIT lab, Toulouse, France), and a first-year master’s program in data science (UT1 Capitole, Toulouse, France).

To minimize misunderstandings, a training session of about 15 minutes was held prior to the study to explain the objective of the study, present the definition of *discernibility* as described above, the interface to receive the information and the various graphics available without influencing the subsequent study.

### 2.3. Data sources

Data originate both from human-based dataset and synthetic generation.

#### 2.3.1. Human-based dataset

In order to have a dataset of sufficient size, with variables collected in humans and whose significance is familiar to a biomedical audience, a subset of data coming from The National Health and Nutrition Examination Survey in the United States (NHANES) has been chosen. As previously detailed [16], the dataset consists of computing physiological age from 60,402 individuals and 48 biological variables. Subsets of variables were randomly generated and used to train ML models. SHAP was selected as the XAI method for its model-agnostic and additive nature [17]. By varying both the type of ML model and the subsets of variables, 11,872 explanations were generated, each one representing a single variable in a subset with a data value vector and a SHAP influence value vector. The aim is to achieve maximum diversity in the nature of the relationship between variable values and explanations for a given variable. Therefore, the statistical coefficients (Pearson correlation coefficient, Spearman’s *ρ* coefficient, Kendall’s *τ*, cosine correlation, distance correlation, xi correlation) between the data value and SHAP influence value were calculated to describe the characteristics of each of the 11,872 explanations. Using K-Means clustering optimized by achieving the best possible silhouette scores, 15 representative explanations were ultimately identified and evaluated in this study.

#### 2.3.2. Synthetic data sources

Since some patterns of relationship were absent from the human-based dataset such as linear or periodic relationships, the synthetic explanations were generated using mathematical functions that encompassed both monotonic and non-monotonic, linear and non-linear distributions. Additionally, data noise was introduced to evaluate the impact when the values of features have varying or non-corresponding influence values, which is a critical factor affecting explanation quality. The following functions were used, with *x* the raw values of a variable, and *y* the related values of explanations (SHAP values): linear function (*y* = 0.5 *× x*), polynomial function (*y* = *x*^2^), root function 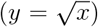, trigonometric functions (*y* = *sin*(*x*), *y* = *arcsin*(*x*)). Composite functions were not considered, being combinations of main functions. A total of 10 synthetic explanations were considered (5 without and 5 with added noise).

#### 2.3.3. Conduct of the study

During the experiments, participants were first asked to evaluate synthetic explanations, followed by NHANES explanations. The explanations were presented in two visual formats: 1) A summary plot alone (a scatter plot showing influences for each variable and each individual with raw data as color code), and 2) a combination of a summary plot and a partial dependence plot (a scatter plot showing the relationship between data and explanations for each variable and each individual).

For each type of explanation, the assessment involved first evaluating the summary plot in a randomly determined order and subsequently assessing the combination plots in another randomly determined sequence. Participants rated discernibility on a scale from 0 to 100 and provided a confidence level for the score (1 to 5, the lowest to highest confidence, respectively). Representative examples of the assessed combination plots are presented in Fig. 1.

**Figure 1.**
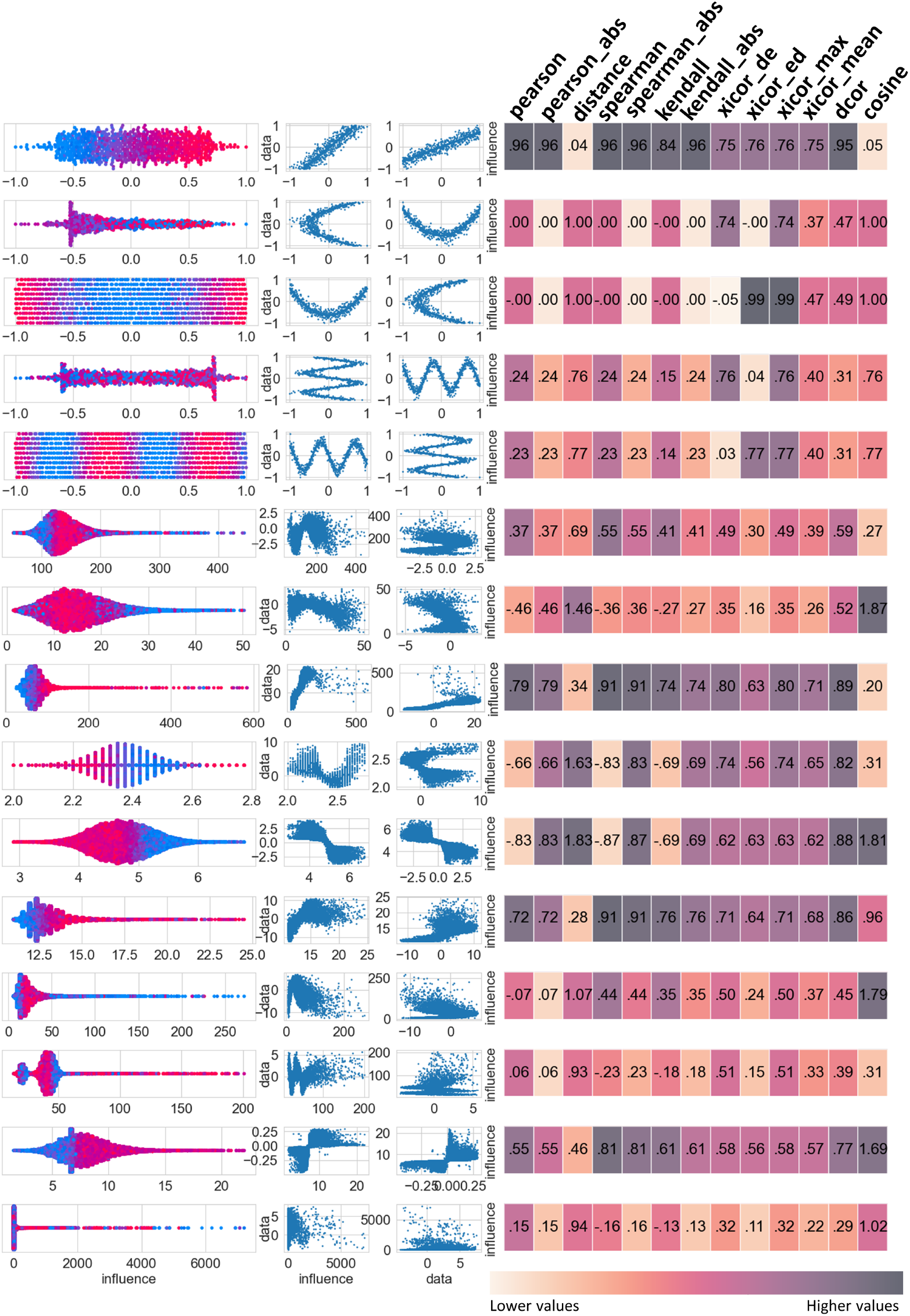
Examples of combination plots. The first 5 plots were generated synthetically, while the remaining plots sourced from NHANES datasets. The panel incorporates the combination of the summary plot on the left, followed by the partial dependence plot. Statistical coefficients are provided beside each respective plot, the color indicates the normalized value of a given coefficient.

#### 2.3.4. Statistical coefficients as candidates for discernibility

The aim of these statistical coefficients is to be able to capture for a given variable the relationship between its values and its explanations. The first candidates were based on monotonicity, specifically Spearman’s *ρ* (*spearman*) and Kendall’s *τ* (*kendall*). In addition, Pearson’s correlation coefficient (*pearson*), which assesses linear correlation through covariance, and its transformation, distance correlation (*distance*), were included as candidate. Given that the direction of correlation may be not informative, the absolute value was also applied to the three aforementioned coefficients (*abs*). Additionally, cosine distance (*cosine*) between the data value and SHAP influence value was also calculated following Min-Max normalization.

Because of the expectation of recognizing non-monotonic relations, other coefficients were added as candidates. For example, the distance correlation (*dcor*) [18], which maps the data to a high-dimensional space with a “double centering” step, and xi correlation (*xicor*) [19], a coefficient of correlation designed based on the idea of ties. Due to the asymmetry of the xicor, both directions of dependence were calculated as candidates, hence obtaining xicor for variable/explanation (i.e., *xicor_ed*) and explanation/variable (i.e., *xicor_de*) relationships. Their mean and maximum value are also considered (i.e., *xicor_mean* and *xicor_max*).

We propose a weighting based on their proportion of SHAP influence using Formula 1, where *inf* (*i*) signifies the influence of the feature *i*^*th*^, calculated by averaging the absolute values of the influence across all instances for the *i*^*th*^ feature, and *disc*(*i*) denotes the discernibility of the *i*^*th*^ feature, computed using *dcor*.

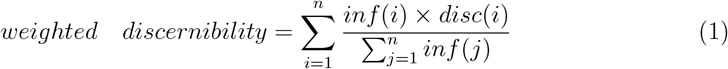

### 2.4. Statistical analyses

The initial part of investigation aimed to determine internal consistency within various categories based on participants’ academic backgrounds and their knowledge of XAI. The intraclass correlation coefficient (ICC) [20] calculates the reproducibility among independent observers with the “irr” R package. The ICC was interpreted as excellent reliability when the value is greater than 0.9, good if greater than 0.75, moderate if greater than 0.5 [21].

The Passing-Bablok regression [22] was employed to assess the agreement between the ratings provided by each individual participant and the tested statistical coefficients. All scores were normalized to the range 0-1. Hence, for each subject of the study, the Passing-Bablok regression provided the value of the slope with an intercept and a 95% confidence interval (95%CI) for both slope and intercept. This study prioritizes the slope value and the width of its 95% CI given that the systemic error (intercept) is common in human evaluation [23]. Bland-Altman plots [24] were drawn to analyze the distribution of residuals with a potential differential bias between high and low values.

## 3. Result

A total of 50 responses were obtained, 19 from participants with biological backgrounds (physiologists, physicians) and 31 with data science background.

### Users’ perception of the discernibility of explanations is improved by partial dependence plots, but remains poorly reliable among users

The ICC scores indicated that the inter-rater reliability was poor (*<* 0.5) whatever level of knowledge or field of expertise, whether assessed through summary plots alone or in combination plots with partial dependence plots (Table 1). Interestingly, providing dependence plot significantly increases users’ confidence in the evaluation of discernibility (Table 1). Confidence was significantly improved with level of education (mean improvement of confidence between combination plots and summary plots of 0.02, 0.23 and 0.36 for Bachelor, Master and Doctoral degrees, respectively), for biomedicals but not data scientists (mean improvement of 0.31 and 0.1, respectively) and for users unfamiliar with XAI outputs compared to those familiar with XAI outputs (mean improve of 0.23 and 0.14, respectively). An increase in confidence comes with a drop in reliability (Table 1). Knowing that the two types of plots provided identical information albeit in different visual formats, this discrepancy may reflect an understanding bias due to different interpretations of visualizations, potentially hindering collective consensus. For this reason, an objective evaluation, such as a numeric value, could aid users in capturing discernibility and assist them in comprehending model explanations.

**Table 1:** Reliability of user’s perception of discernibility of explanations. The intraclass correlation coefficients (ICC) with 95% confidence interval are provided for both summary plots and combination plots, stratified according the type of plots, type of subject’s expertise, their knowledge about explainable artificial intelligence (XAI) and their level of education. p-value relates to a possible significant difference in confidence level between summary plots only and combination plots.

|  |  | N | Summary plots only |  | With combination plots |  | p-value of |
| --- | --- | --- | --- | --- | --- | --- | --- |
|  |  |  | ICC [95% CI] | Confidence | ICC [95% CI] | Confidence | confidence difference |
|  | **All** | 50 | 0.44 [0.31; 0.62] | 3.32 ± 1.08 | 0.35 [0.24; 0.53] | 3.49 ± 1.06 | < 0.001 |
|  | **Type of plots** |  |  |  |  |  |  |
|  | Synthetic | 50 | 0.52 [0.34; 0.77] | 3.38 ± 1.14 | 0.35 [0.20; 0.63] | 3.66 ± 1.07 | < 0.001 |
|  | NHANES | 50 | 0.36 [0.20; 0.64] | 3.25 ± 1.01 | 0.37 [0.22; 0.65] | 3.32 ± 1.02 | 0.1845 |
|  | **Type of expertise** |  |  |  |  |  |  |
|  | Biologist | 19 | 0.45 [0.31; 0.64] | 3.37 ± 1.04 | 0.35 [0.22; 0.53] | 3.68 ± 0.96 | < 0.001 |
|  | Data scientist | 31 | 0.43 [0.30; 0.61] | 3.28 ± 1.10 | 0.36 [0.24; 0.54] | 3.38 ± 1.11 | 0.1229 |
|  | **Knowledge about XAI** |  |  |  |  |  |  |
|  | Rather unfamiliar | 19 | 0.52 [0.38; 0.70] | 3.28 ± 1.05 | 0.32 [0.20; 0.51] | 3.51 ± 1.06 | < 0.001 |
|  | Rather familiar | 31 | 0.41 [0.28; 0.59] | 3.34 ± 1.09 | 0.37 [0.25; 0.55] | 3.48 ± 1.07 | 0.0193 |
|  | **Level of education** |  |  |  |  |  |  |
|  | Bachelor’s degree | 20 | 0.38 [0.25; 0.57] | 3.33 ± 1.09 | 0.39 [0.26; 0.58] | 3.35 ± 1.10 | 0.73 |
|  | Master’s degree | 19 | 0.49 [0.35; 0.67] | 3.34 ± 1.02 | 0.36 [0.23; 0.55] | 3.57 ± 0.98 | 0.0012 |
|  | Doctoral degree | 11 | 0.46 [0.31; 0.66] | 3.25 ± 1.16 | 0.31 [0.18; 0.51] | 3.61 ± 1.11 | < 0.001 |

### dcor, a statistical metric capable of capturing the intra-individual reliability of discernibility perceived by users

Given that there is no consensus of discernibility between users, a Passing-Bablok regression was performed for each user to highlight the consistency between user rating and each statistical coefficient candidate. As revealed by a median slope closer to 1 and a narrower confidence interval width for the Passing-Bablok regression, *xicor ed, spearman abs, pearson abs, kendall abs* and *dcor* coefficients appear suitable candidates for discernibility (Fig. 2A and B). Bland and Altman plots show that *dcor* is the statistical coefficient with the lowest differential bias between the most and least discernible values (Fig. 2). As a result, *dcor* was chosen as the statistical coefficient that better reflects the user’s perception of discernibility.

**Figure 2.**
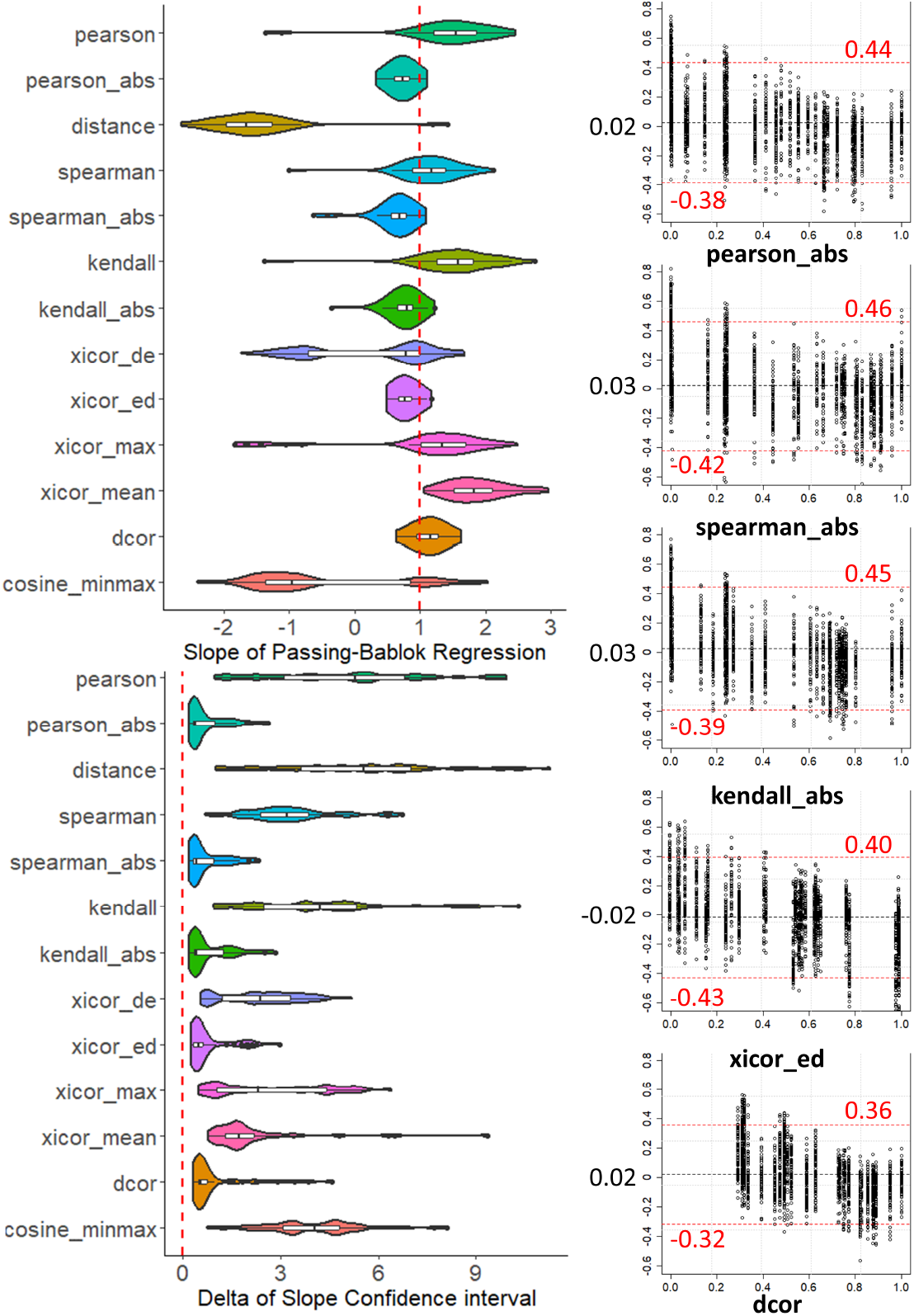
Ability of the *dcor* metric to better reflect the human perception of discernibility of the explanations. In order to capture the relationship between the coefficient and user perception by integrating reliability at the individual user level, a Passing-Bablok regression has been performed. The two violin plots depict the distributions of the slope and its confidence interval width obtained from the Passing-Bablok regressions performed for each user. On the right, Bland-Altman plots illustrate the residuals of each statistical coefficient on the y-axis against the coefficient values on the x-axis. The red dashed lines indicate the 95% confidence intervals.

### Using dcor for discernibility as an additional metric to accuracy for obtaining more acceptable ML models

Since the more influential variables were supposed to be more discernible, the weighted discernibility has been used (Formula 1). Weighted discernibility allows comparison between different models and feature subsets, complementing traditional evaluation performance metrics for feature selection. Hundred random subsets comprising 20 features from the NHANES dataset were trained on 3 XGBoost models (tree-based models) and 6 artificial neural networks (ANN) with different structures and hyperparameters to predict biological age (regression task). The trade-off between the overfitting rate, the *R*^2^ values, and the weighted discernibility are depicted in Fig. 3. A notable inverse correlation between overfitting and discernibility is observed in the three XGBoost models. Similar patterns are seen in ANN models. The discernibility is able to identify the overfitting model (Fig. 3A plot 1), even if their *R*^2^ values are similar (Fig. 3A plot 2 and 3). The model complexity also leads to a decrease in discernibility as demonstrated in Fig. 3B. Generally, XGBoost exhibits better discernibility compared to ANN. Moreover, increasing the depth of ANN results in a decrease in discernibility without a significant increase in performance.

**Figure 3.**
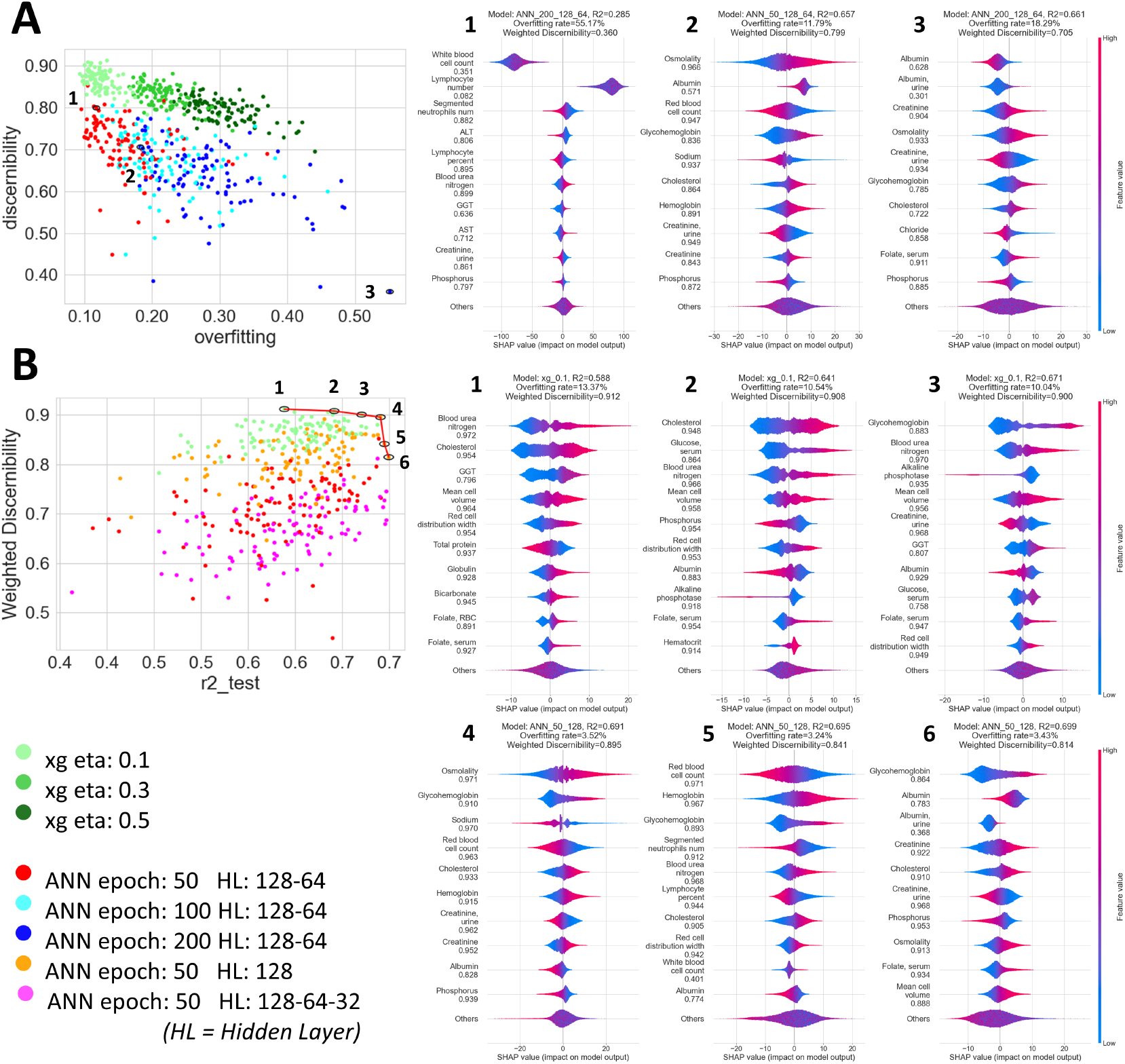
The dcor metric is suitable for feature selection and may prevent ML model overfitting. Model performance and explanation *discernibility* of 100 random subsets comprising 20 variables from the NHANES dataset are presented. Part A illustrates the trade-off between model overfitting (relative change of the *R*^2^ between the train and test dataset) and weighted *discernibility* with the illustration of 3 chosen summary plots (1 to 3). Part B illustrates the trade-off between model performance (*R*^2^ on the test dataset) and weighted *discernibility* with the respective summary plots for each model on the Pareto’s front (1 to 6). For each summary plot, the variables are sorted in descending order of their contribution to the model. For each variable, a dot represents an instance of the dataset (red for a high feature value and blue for a low value). A positive value on the x-axis indicates feature contributes positively to the prediction for this instance, and conversely for a negative value. To enhance the readability, only the top ten most important features were shown.

A Pareto Front can be drawn, composed of six optimal solutions of feature selection for the NHANES dataset (Fig. 3B). Each solution (Fig. 3B plot 1 to 6) can be interpreted as a trade-off between performance and discernibility. For example, users can choose between subsets *4* and *5*, which respectively achieved a better *discernibility* (0.895 for subset *4* and 0.841 for subset *5*) or *R*^2^ (subset *5*). If the problem of feature selection expands to model selection, the solution of subset *3* with the XGBoost model reaches a higher *discernibility* (0.90) with a slight decrease of *R*^2^ (0.67).

## 4. Discussion

The distance correlation (*dcor*) has been shown to be a reliable method for assessing the *discernibility* of explanations, effectively capturing the clarity of the relationship between data and the explanations. Additionally, this metric offers an objective means of selecting variables in ML models whose explanations would be more acceptable to health-care professionals without impairing model performance and preventing overfitting.

In current XAI evaluations, significant emphasis is placed on application-grounded assessments, notably *plausibility* [12] *and user satisfaction* [25]. These evaluations typically require a case-by-case examination by domain experts, drawing upon their existing knowledge and experience. However, this approach is both resource-intensive and impractical for automated calculation. Similar issues occur in other metrics, such as *interestingness* [26] *and informativeness* [27], which require less domain-specific knowledge, but still include subjective measures such as human judgment (e.g., assessing *surprisingness* and usefulness), rendering them unsuitable for automated computation. Conversely, *discernibility* seeks to uncover the explorable relationships within explanations using a statistical coefficient, *dcor* which is the best fitting when considering the subjects/user’s level. Compared to *readability* [13], its notable advantage lies in accommodating non-monotonic relationships, which is indispensable in biology.

When discussing relationships, it’s tempting to revisit the classic fallacy concerning correlation and causality, which was first identified in statistical analysis [28] and extended later to the ML analysis. Correlation does not imply causation. In ML models, correlation can stem from causation, confounding factors, or selection bias [29], the latter two are termed spurious correlations because they do not necessarily indicate causality in the model. In this context, a discernible feature may be interpreted as a high correlation but should not be assumed to imply causality from the feature to the target. Causal inference may be derived from the model’s explanation but requires additional causal computation, such as Bayesian causal networks [30]. Although this fallacy is well recognized in the ML community, it may not be as obvious to end-users [31]. Implementing *discernibility* in an XAI-based decision support system should include clear explanations of these fundamental terms to help users avoid this potential misunderstanding. However, the nature of the relationship between variables and their influence may provide clues to the underlying biological mechanisms that are not immediately apparent when examining individual predictors.

The consideration of user profiles is also essential for designing systems that meet their needs. For example, the additional dependency plots may be more beneficial for biologists than data analysts in improving users’ confidence. Besides, confidence in the system is reinforced by *discernibility*, which ensures meaningful explanations are provided, as required by *Four Principles of Explainable Artificial Intelligence*^3^. Such as in healthcare, groups of patients can be identified by shared explanations [32], and the discernible relationship, or the patterns found, would be much more informative to the biomedical expert and open up the possibility of obtaining causal insight. Compared to other similar expert-based studies (in general less than 10 [33, 34]), the current study achieved a sizable number of participants. Moreover, the *discernibility* overcomes the bias due to the educational background or personal experience and provides an objective reference in the understanding of explanation. This brought the potential of assisting users in other research fields, such as finance, ecology, automobile, etc.

Another potential application of *discernibility* lies in model verification. Low *discernibility* may indicate the need to revise the model’s hyperparameters to prevent overfitting. Additionally, the complexity of ML algorithm plays a crucial role. While complex models often offer superior predictive capacity, their reasoning process can be opaque, hence difficult to be understood by humans. XAI was developed to address this problem [4], yet the explanatory capacity varies still across the model complexity. As *discernibility* illustrated, the clarity of explanation declines as model complexity increases. Therefore, users should be empowered to make informed decisions, balancing the trade-off between model complexity and explainability according their individual needs.

Feature selection also faces similar issues. With the development of clinical examination technology, the number of indicators available for disease diagnosis and prevention has increased, including probably redundant and useless parameters. Hence, FS has become an essential step of the biomedical ML pipelines [35]. Traditional FS methods typically rely on model performance metrics (e.g., accuracy, *R*^2^) to identify the most suitable subset [36]. However, metrics based on XAI could complement these traditional methods by helping to identify subsets that retain as much information as possible [37]. The concept of *discernibility* enables the selection of feature subsets that display clear relationships, which is desirable in the sense-making of the model. In this multi-objective optimization, a set of solutions could be proposed to the users, enabling users to choose according to their preferences. Although *discernibility* is helpful in providing informative explanations, it cannot replace humans in decision-making. The interaction between humans and the XAI system (*i*.*e*., interactive XAI) brought the value added in supporting the human cognitive process of explaining [38]. According to a human-in-the loop approach, users should either express their strong preference for a particular goal of the FS (e.g. maximize accuracy, minimize the difference in explanations with the model without FS, maximize discernibility of explanations) [37] and thus have the possibility of visually exploring the different optimal alternatives, judging them based on their own domain knowledge.

### 4.1. Conclusion

The *discernibility* approach can help meet today’s challenges in XAI and the biomedical field since it has the strength to provide an objective reference metric which helps users understand the model regardless of their personal background and knowledge. It can help better identify explanations that make sense to the biomedical expert (and avoid analyzing explanations that have little relationship with the original data), thus achieving more acceptable explanations and ML models. It is essential to incorporate all potential dimensions (e.g., accuracy, retention rate, similarity to the original explanation, discernibility) into a comprehensive process. A conclusive decision could then be suggested for feature selection based on user priorities or a trade-off determined by optimization algorithms for decision-making.

## 5. Acknowledgments

This work was supported by the Occitanie Region, the Federal University of Toulouse Midi-Pyrénées (grant ADI 2021, N°ALDOCT89533). This study has been partially supported through the grant EUR CARe N°ANR-18-EURE-0003 in the framework of the Programme des Investissements d’Avenir and the national infrastructure “ECELLFrance: Development of mesenchymal stem cell based therapies” (PIA-ANR-11-INBS-005).

## Footnotes

1 https://eur-lex.europa.eu/legal-content/EN/TXT/?uri=celex%3A52021PC0206

2 https://www.whitehouse.gov/briefing-room/presidential-actions/2023/10/30/executive-order-on-the-safe-secure-and-trustworthy-development-and-use-of-artificial\protect\discretionary{\char\hyphenchar\font}{}{}-intelligence/

3 https://www.nist.gov/publications/four-principles-explainable-artificial-intelligence

